# Rapid integration of somatosensory feedback in planning tongue movements

**DOI:** 10.64898/2026.03.03.709371

**Authors:** Jeong Jun Kim, Mingyuan Dong, Daniel H. O’Connor

## Abstract

The brain dynamically generates motor actions based on fast-changing sensory feedback arising from previous actions and the environment.^1,2^ How neural circuits carry out flexible motor control at fast timescales remains incompletely understood. Here we developed a probabilistic sequence licking task, in which mice adjust lick angles from lick to lick based on lingual somatosensory feedback. Motor planning in sequences of licks flexibly directed on-the-fly depended on orofacial cortical activity on a lick-by-lick timescale. Population activity in the orofacial cortices and superior colliculus generates an internal estimate of lick target position, based on cortical processing of somatosensory feedback. Persistent memory of lick target position absent sensory feedback is maintained in the orofacial motor cortex. Our results show how a multiregional circuit carries out sensory processing, motor preparation, and motor execution within rapid sequences of orofacial movements.

## Introduction

The brain continually transforms sensory feedback into motor actions, which in turn generate new sensations based on the interactions between motor actions and the environment.^1,2^ An intriguing and unresolved question is how the brain continually plans and adjusts motor actions moment to moment based on new sensory feedback.

Motor planning and execution have been studied extensively in delayed response tasks that separate sensory cue and eventual motor action in time. Studies in reaching tasks and directed licking tasks have shown how population activity in cortical and subcortical areas encodes the flow of information from sensory processing to motor preparation and execution of a single well-prepared movement.^3–11^ However, how sensory processing, motor preparation, and motor execution occur during sequences of movements flexibly constructed on-the-fly remains incompletely understood. Execution of current movements and planning of future movements based on current sensory feedback can occur rapidly and at every step of a sequence,^12–14^ and neural mechanisms responsible for such computations remain to be elucidated.

Here we show that mice redirect the licks in a motor sequence from lick to lick based on somatosensory feedback at the tongue. Planning the direction of the lick depends on orofacial cortical activity on a lick-by-lick timescale. We then demonstrate that the population activity in orofacial cortex and superior colliculus compute an internal estimate of lick target position, which relies on cortical processing of somatosensory feedback. Persistent memory of lick target position is maintained in the orofacial cortex. Together, we show how a multiregional circuit for lingual control processes sensory feedback, plans future movements, and executes current movements on the fly.

## Results

### Mice adjust lick angles from lick to lick based on lick contact

To investigate how licks are planned and executed moment to moment, we developed a head-fixed sequence licking task with unpredictable somatosensory feedback (Fig. 1a, Supplementary Video 1). We trained water-restricted mice to lick at a motorized lick port for water with the following trial structure (Fig. 1c): following an auditory cue at the start, the position of the lick port would shift either right or left with equal probability (i.e. random walk) after every third lick contact with the lick port.

**Figure 1.**
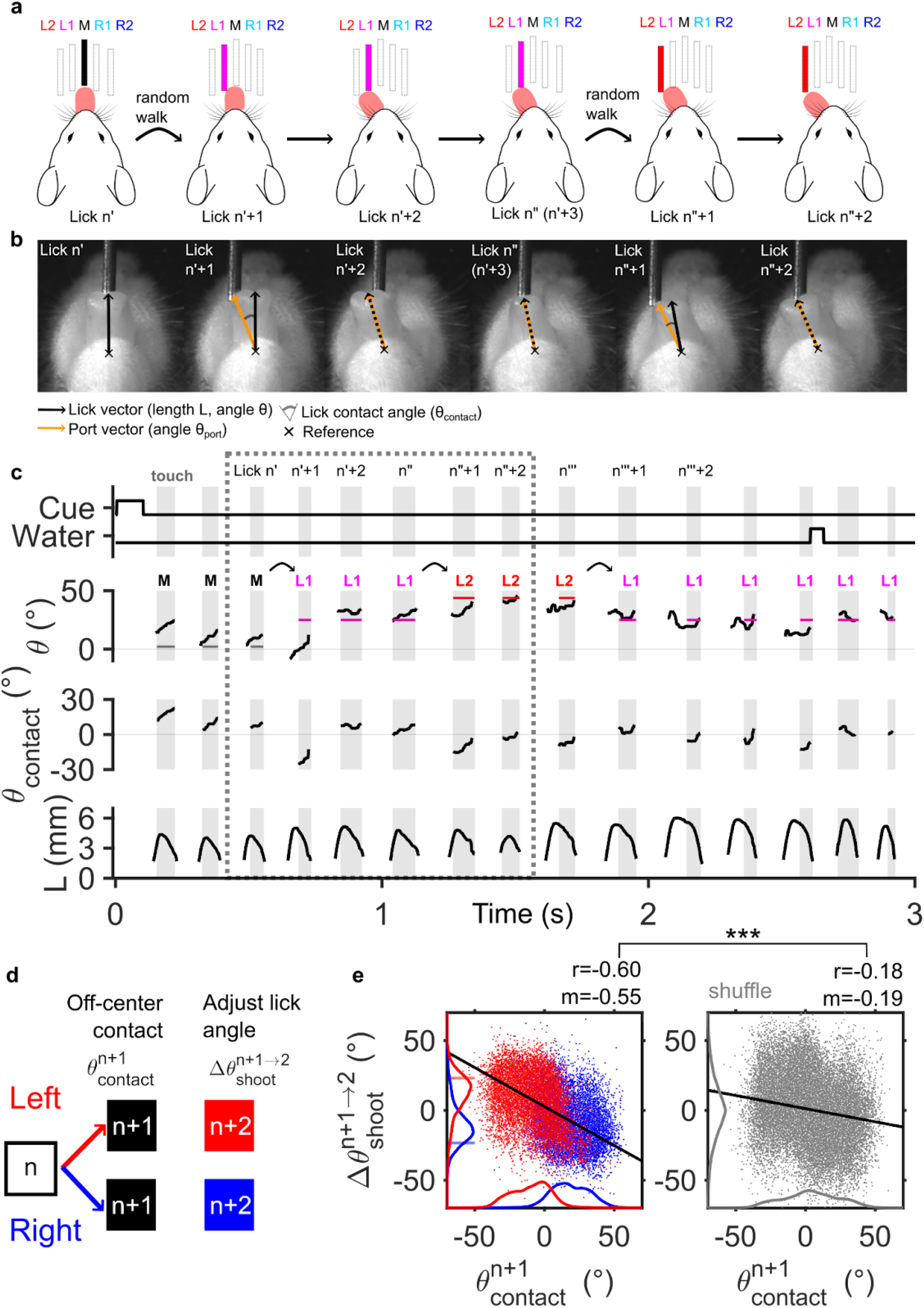
Probabilistic sequence licking task. **a**. Schematic of the probabilistic sequence licking task. A motorized lick port undergoes a random walk between L2 (leftmost), L1, M, R1, and R2 (rightmost) lick port positions (colored red, pink, gray, cyan, blue), shifting left or right from the current position with equal probability. Licks are labeled relative to the timing of the random walk, such that lick n (n’,n”,n’’’) is the lick immediately prior to the random walk, lick n+1 (n’+1,n”+1,n’’’+1) is the first lick after the random walk, and similarly for lick n+2. **b**. Bottom-view still frames for a series of licks (n’, n’+1, n’+2, n”, n”+1, n”+2) in an example trial, corresponding to licks in the dotted box in **c**. Lick vector of length L and angle θ is defined as the vector between a fixed reference point and tongue tip. Port angle θ_port_ is defined as the angle of the vector between the reference point and midline of the lick port tip. Tongue contact angle is defined as the difference between lick angle θ and port angle θ_port_. **c**. Time series of task events and behavioural variables during an example trial. Periods of tongue–port contact are shaded in grey. Curved arrows indicate random walks between port positions. Still frames in **b** are from licks in the dotted box. Port angle θ_port_ is overlaid and labelled in the plot of lick angle θ. **d**. Diagram of lick angle aiming for lick n+2 following contact angle at lick n+1 following a random walk. Off-center contact occurs at lick n+1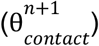, which leads to adjustment in lick shooting angle in the subsequent lick 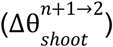. **e**. (Left) Scatter plot of pairs of the lick n+1 contact angle 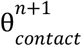 and change in lick shooting angle 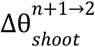 (n= 40316 lick pairs, 72 sessions, 7 mice). Kernel density estimations of each variable are overlaid on the axes. Right and left random walks are color coded blue and red, respectively. Horizontal bars on y-axis indicate the average change in lick port angle θ_port_ during right and left random walks (blue and red). Black line indicates the line of best fit from linear regression. Pearson’s correlation coefficient r and slope of linear regression m shown in top right. (Right) Scatter plot and kernel density estimation for shuffled pairs of lick n+1 contact angle 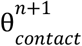 and subsequent change in lick shooting angle 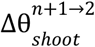; same dataset as left but shuffled. *** p<0.001, one-sided shuffle test.

We denote the lick immediately prior to random walk of the lick port as lick n, the first lick following random walk as lick n+1, and so forth for lick n+2 and n+3. The third lick contact with the lick port triggered a new random walk. The lick contact with the lick port was detected in real-time and used to drive trial progression.

A trial consisted of three random walks and a droplet of water reward at the end. After a random inter-trial interval, the next trial would begin from the final position of the lick port at the end of the previous trial. High speed videography at 400Hz frame rate and markerless pose estimation^15^ were used to track the position of the tongue tip (Methods), which was used to compute the length and angle of a given lick (Fig. 1b).

Given the auditory masking conditions and absence of visual cues, the most salient sensory feedback for the task was lingual somatosensation, specifically the senses of touch and proprioception for the tongue. The displacement of the lick port in a random walk, approximately equal in distance to half the tongue width, caused off-center contact that indicated the new position of the lick port after a random walk. The off-center contacts were quantified with lick contact angle (θ_contact_), the difference in the angle between the tongue tip (θ) and tip of the lick port (θ_port_) (Fig. 1b).

In the probabilistic sequence licking task, mice continually adjusted the lick angle to follow the random walks of the lick port. In the first lick after a given random walk (lick n+1), the tongue typically encounters an off-center contact event (large |θ_contact_|). In the subsequent lick n+2, the lick angle adjusts to re-center the lick port to the tongue midline (decreased |θ_contact_|) (Fig. 1d), a strategy that minimizes the number of licks that miss the lick port during the sequence. The adjustment in angle between the first and second lick was quantified as the difference in the shooting angle in each lick or 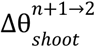 (Methods). On a lick-by-lick basis, the lick angle adjustment between lick n+1 and n+2 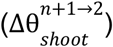 was anti-correlated (r = −0.60) and had a negative linear relationship (slope m = −0.55) with the lick n+1 contact angle 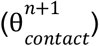

well-above chance levels (p<0.001) (Fig. 1e). Anti-correlated and negative linear relationships above chance levels held true for each individual session and for each animal in the dataset (Extended Data Fig. 1). The anti-correlated and negative linear relationship held true for the subsequent pair of licks n+2 and n+3 (Extended Data Fig. 5c), suggesting that lick targeting based on somatosensory feedback from the previous lick cycle occurs at every step of the lick sequence.

### Motor planning in probabilistic sequences depends on cortical activity on a lick-by-lick timescale

Orofacial cortical areas, including anterolateral motor cortex (ALM), tongue-jaw primary motor cortex (M1TJ), and tongue-jaw primary somatosensory cortex (S1TJ), are known to be critical in motor and sensory processing in various licking tasks.^3,16–19^ To test whether executing probabilistic sequences relies on orofacial cortical areas, we used Vgat-Cre/Ai32 mice to bilaterally inhibit orofacial motor cortex (ALM/M1TJ) or somatosensory cortex (S1TJ) (Extended Data Fig. 2). Given the proximity of ALM and M1TJ (~1.3 mm), the motor cortical regions were grouped together for the purposes of photoinhibition. Consistent with previous studies,^3,17,18^ bilateral ALM/M1TJ photoinhibition over multiple lick cycles (2 s) strongly suppressed lick production and shortened lick lengths, while bilateral S1TJ photoinhibition increased variability in lick angles (Extended Data Fig. 2c-e, Supplementary Video 2, 3). Optogenetic silencing of barrel field cortex (S1BF), a cortical area not associated with lingual somatosensory or motor functions, did not decrease lick rates, length, or accuracy in lick angles (Extended Data Fig. 2c-e), demonstrating nonspecific cortical silencing was not responsible for the observed changes in lick kinematics.

We set out to perturb cortical activity associated with individual licks, particularly lick n+1; the off-center contact at lick n+1 would result in somatosensory signals and motor planning to redirect the subsequent lick n+2. Using closed loop optogenetics (Methods), we photoinhibited ALM/M1TJ and S1TJ over lick cycle n+1 (Fig. 2, Extended Data Fig. 3). We showed that brief optogenetic perturbation in the orofacial cortex disrupts subsequent lick targeting (Supplementary Video 4, 5). For both ALM/M1TJ and S1TJ photoinhibition experiments, cortical silencing significantly reduced the anti-correlation between lick n+1 contact angle and lick angle adjustment between lick n+1 and n+2 (p<0.001) (Fig. 2b,c). Instead, the tongue “revisited” in lick n+2 the same angle as lick n+1 with minimal change in shooting angle 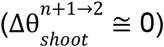, indicating failure to update upcoming motor action based on somatosensory feedback.

**Figure 2.**
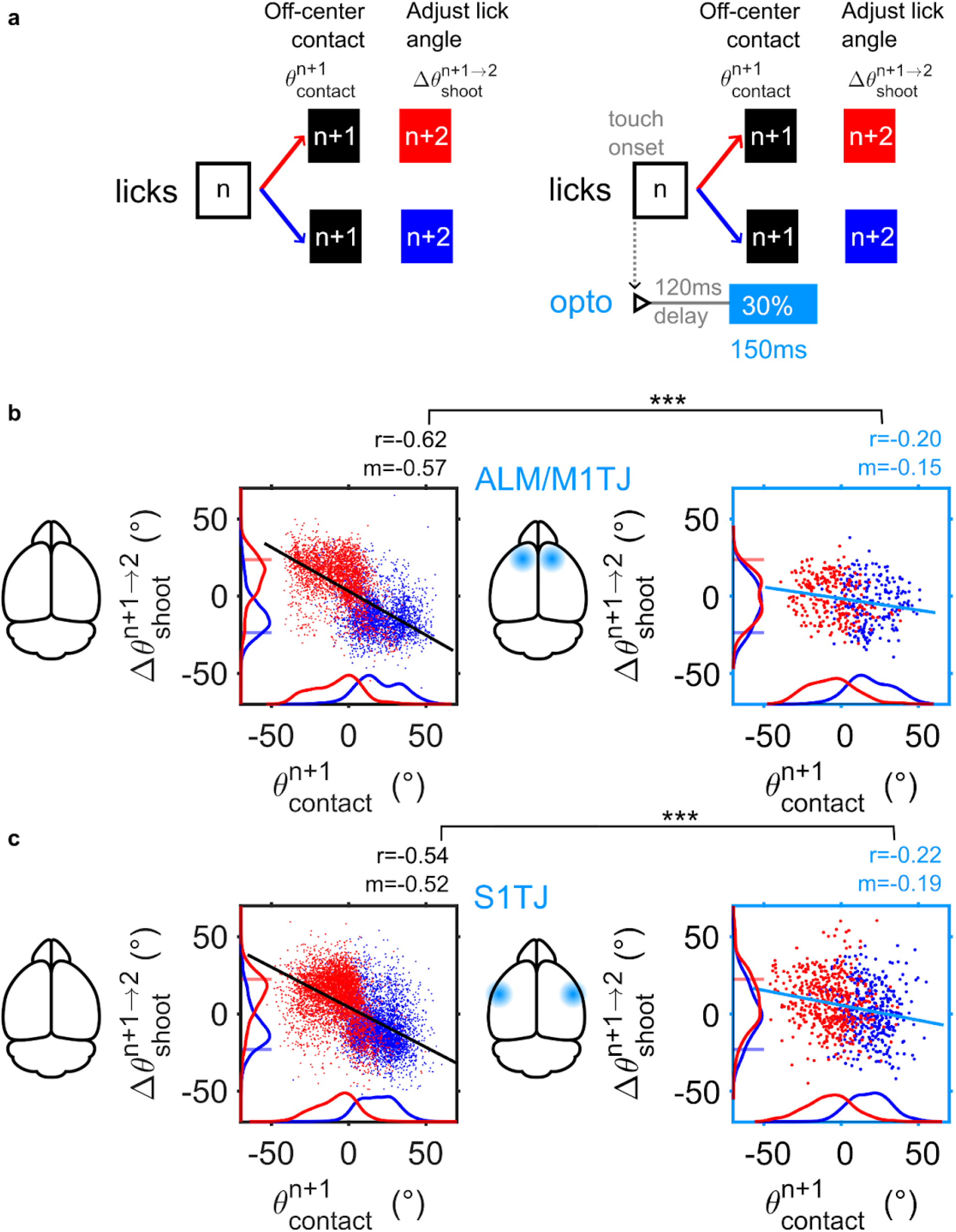
Motor planning in probabilistic sequences depends on cortical activity on a lick-by-lick timescale. **a**. Diagram of closed loop optogenetic perturbation of lick n+1. Photoinhibition (40Hz sinusoid with ramp-down) occurs randomly in 30% of trials following a 120 ms delay after touch onset of lick n and lasts for 150 ms (Methods). **b**. Scatter plot and kernel density estimation of lick n+1 contact angle 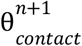 with or without ALM/M1TJ photoinhibition (right, left) and subsequent change in lick shooting angle 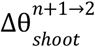(n= 5433 control lick pairs, 464 opto lick pairs, 10 sessions, 3 mice). Colors as in Fig. 1d. ALM, anterolateral motor cortex; M1TJ, tongue-jaw primary motor cortex; S1TJ, tongue-jaw primary somatosensory cortex. *** p<0.001, one-sided paired shuffle test. **c**. Scatter plot and kernel density estimation of lick n+1 contact angle 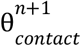 with or without S1TJ photoinhibition (right, left) and subsequent change in lick shooting angle 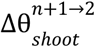 (n= 8748 control lick pairs, 868 opto lick pairs, 17 sessions, 4 mice). Colors as in Fig. 2b.

Notably, the shooting angle of lick n+1 itself was not affected by cortical photoinhibition during lick cycle n+1 (Extended Data Fig. 4h, j), and variability in photoinhibition timing within a lick cycle did not affect lick angle for lick n+1 or lick n+2 (Extended Data Fig. 4i, k). Variability in photoinhibition timing did affect lick length, such that photoinhibiting the early phase of a lick cycle reduced lick length in a graded manner (Extended Data Fig. 4m, o). This observation suggests that cortical control of lick length operates on an intralick timescale. In contrast, lick angle targeting depended on cortical activity on a lick-by-lick timescale, not within-lick cycle timescale. Silencing orofacial cortical activity disrupted planning the next lick angle, but not execution of the current lick angle.

Photoinhibiting motor cortex during lick cycle n+2 caused similar impairment in lick angle adjustments between licks n+2 and n+3 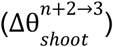 (Extended Data Fig. 5c), showing that regardless of which lick cycle was targeted by optogenetic silencing, the subsequent lick had impaired angle targeting. The optogenetic perturbation of lick targeting suggested that the cortical activity during the current lick cycle contains the feedback and control signals necessary for planning the subsequent lick.

### Population coding of lick target in orofacial sensorimotor cortex and superior colliculus

The probabilistic sequence licking task requires the brain to track the lick target location, inferred through somatosensory feedback in contact events, and update its lick angle accordingly. Here we propose a putative algorithm for the task (Fig. 3a), in which proprioceptive sensing of current lick angle (θ) and tactile feedback from lick contact (θ_contact_) is combined to generate an internal estimate of the port angle (θ_port_). The estimated port angle is then used to update the intended angle (θ) for the upcoming lick, and the process is repeated lick-by-lick, with timescale matching the lick cycle (~ 6.5 Hz).

**Figure 3.**
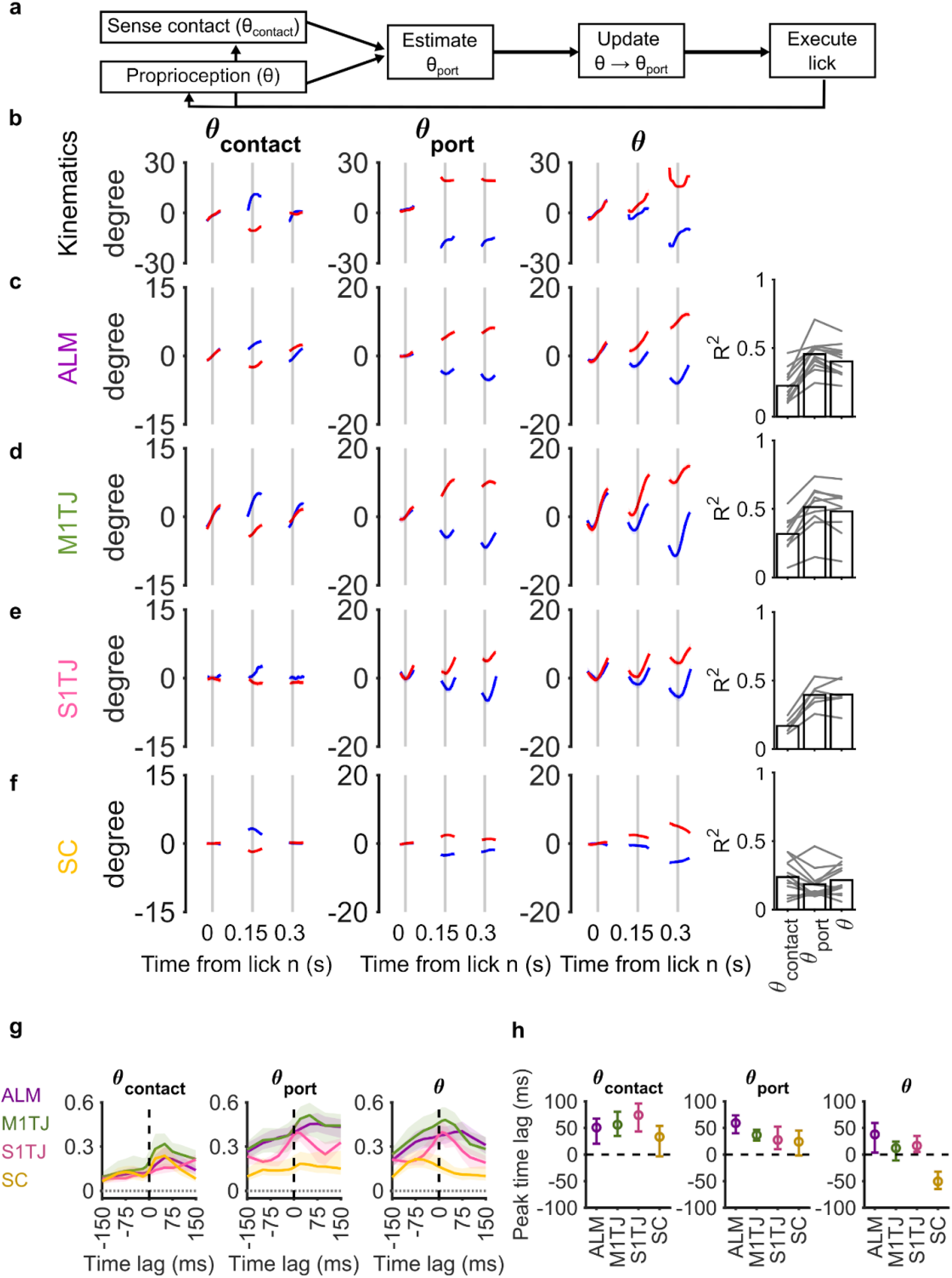
Neural correlates for estimating lick target position are distributed across orofacial cortical areas and superior colliculus. **a**. Proposed algorithm for lick aiming in the probabilistic sequence licking task. At each lick, tactile feedback from tongue-tip contact θ_contact_ and proprioception of current lick angle θ is combined to estimate the angle of the lick port θ_port_ following a random walk. The estimated θ_port_ is used to update the angle of the next lick θ. **b**. Time series of behavioural variables (mean ± 99% bootstrap confidence interval; n = 9663 random walks, 41 sessions) for left (red) and right (blue) random walks. Time series are grouped by random walk direction and trial-averaged. **c-f**. (Left) Linear decoding of behavioural variables (columns) from populations recorded in the ALM, M1TJ, S1TJ, and SC (rows). Displayed traces are results from linear decoding from linear models with appropriate time lags added to spikes; ALM: −50 ms, M1TJ: −50 ms, S1TJ: −20 ms, SC: −50 ms (θ_contact_, θ_port_), +50 ms (θ). Traces are grouped by random walk direction and trial-averaged across multiple sessions (ALM, n = 12 sessions; M1TJ: n = 9 sessions; S1TJ, n = 6 sessions; SC n = 14 sessions). Thicker lines indicate durations of visible tongue, inter-lick intervals omitted. Same plotting conventions as in **b**. (Right) Comparison of mean goodness of fit (coefficient of determination R^2^) of linear regressions of behavioral variables for a given region, ALM, M1TJ, S1TJ, and SC (rows). Goodness of fit from individual sessions in gray. **g**. Goodness of fit (coefficient of determination R^2^) of linear regressions of behavioral variables with a range of time lags in spike times (mean ± 95% bootstrap confidence interval). Positive time lags indicate linear regression with spikes shifted post-event, and vice versa for negative time lags. Black dotted lines indicate linear regression without time lag. Recorded areas are color coded (ALM, magenta; M1TJ; green; S1TJ; pink, SC; gold). Goodness of fit in null models (linear regression with shuffled spike rates; Methods) shown in gray dotted lines. **h**. Comparison of time lag with best goodness of fit (“peak time lag”) for linear regression of behavioral variables (mean ± 95% bootstrap confidence interval). Black dotted lines indicate zero time lag. Recorded areas are color coded (ALM, magenta; M1TJ; green; S1TJ; pink, SC; gold).

To identify neural correlates involved in tracking the lick target, we used Neuropixel probes to record from orofacial cortical areas (ALM, M1TJ, S1TJ) and lateral superior colliculus (SC) (Extended Data Fig. 6). The brain regions were chosen for their known roles in controlling tongue movements.^3,16,17,20,21^ We obtained in total 5623 well-isolated units from 182 recordings. Across all recordings, the estimated median false discovery rate per session estimated based on interspike interval violations^22^ was on average 5.1% across on average 37 units per session (Extended Data Fig. 6b-d, Methods).

For each recording session, we used linear regression on each of the relevant behavioral parameters (θ_contact_, θ_port_, θ) to identify the weighted sum of simultaneously recorded firing rates that best predicted the behavioral parameter (Methods). Using the fitted linear model, the behavioural variables were decoded from population activity on a single-trial basis and grouped by the direction of random walk (Fig. 3c-f, Extended Data Fig. 6f). Population activity in orofacial cortical areas (ALM, M1TJ, and S1TJ) had stronger coding for port angle (θ_port_) than for lick contact angle (θ_contact_) (Fig. 3c-f; right). In contrast, population activity in superior colliculus (SC) had comparable coding for lick contact angle (θ_contact_) and port angle (θ_port_). Coding of lick angle (θ) was present in all the recorded areas.

In addition, we measured cross-validated goodness-of-fit (coefficient of determination R^2^) for linear models fitted with firing rates shifted with a range of time lags spanning a lick cycle to test whether population activity preceded or followed behavioral parameters in time (Fig. 3g). Increased goodness-of-fit with negative time lags, in which population activity preceding in time best predicted a given behavioral parameter, suggested motor activity. Conversely, increased goodness-of-fit with positive time lags with population activity following in time suggested sensory activity.

Linear models for lick angle (θ) and port angle (θ_port_) in orofacial cortical populations (ALM, M1TJ, S1TJ) had broad goodness of fit over a range of time delays (Fig. 3g; middle, right), suggesting both motor and sensory signals present in the neural correlates. In contrast, linear models of lick contact angle in orofacial cortical populations (ALM, M1TJ, S1TJ) and collicular populations (SC) had a one-sided shape with increased goodness of fit for positive time lags (Figure 3g; left), suggesting predominantly sensory signals in the neural correlates.

In comparing time lags with the best goodness-of-fit (“peak time lag”) for each recorded population, we observed that collicular populations (SC) had the smallest time lag for coding lick contact angle (θ_contact_), suggesting low latency in tactile response (Fig. 3h). Coding for port angle (θ_port_) in S1TJ and SC populations preceded ALM and M1TJ, indicating motor areas lagged behind sensory areas in neural correlates for lick target position. Coding for lick angle (θ) in cortical populations had small positive time lags in contrast to the collicular population with negative time lag, indicating mixed motor and proprioceptive signals for lick angle in cortex.

The neural correlates for estimating lick target position were present in orofacial cortical areas and lateral superior colliculus. Representation of somatotopic tactile feedback (θ_contact_), lick target position (θ_port_) and lick angle (θ) were distributed across cortical areas and superior colliculus with differing strengths of coding and time lags in neural correlates.

### Optogenetic perturbation of somatosensory processing disrupts estimation of lick target in motor cortex

How does the photoinhibition of cortical activity affect the neural code for targeting licks accurately? We sought to investigate how perturbation of somatosensory processing affects neural correlates in the motor cortices (ALM, M1TJ). We hypothesized that optogenetic perturbation of somatosensory processing would disrupt the following processes in the motor cortex: (1) the update of the internal estimate of lick port location (θ_port_) based on tactile feedback (θ_contact_) and (2) the update of the intended lick angle (θ) for the subsequent lick based on the current estimate of port location (Fig. 4a).

**Figure 4.**
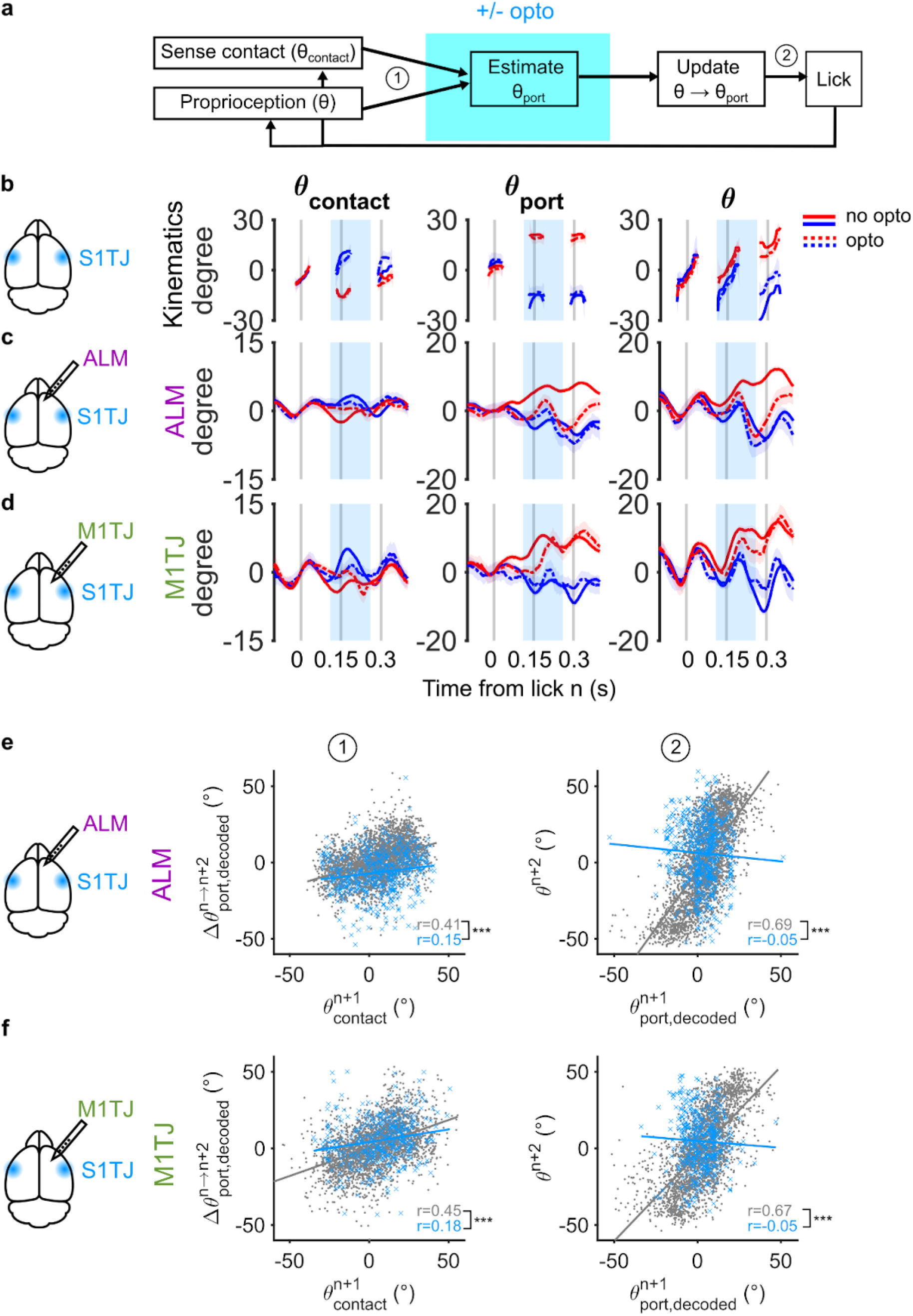
Optogenetic perturbation of somatosensory processing disrupts internal estimation of lick target position by motor cortex. **a**. Optogenetic perturbation of somatosensory processing in the putative lick aiming algorithm. Neural correlates for lick aiming will be perturbed following disrupted somatosensory processing. **b**. Time series of the behavioural variables (mean ± 99% bootstrap confidence interval; n = 6619 random walks without optogenetic perturbation in S1TJ, n = 857 random walks with optogenetic perturbations, 21 sessions) for left (red) and right (blue) random walks. Time series are grouped by random walk direction and trial-averaged. Duration of optogenetic perturbation shown in shaded cyan box. **c-d**. Linear decoding of behavioural variables (columns) from populations recorded in the ALM and M1TJ (rows) with or without optogenetic perturbation in S1TJ (dashed and solid lines, respectively). Displayed traces are results from linear decoding from linear models with appropriate time lags added to spikes; ALM: −50 ms, M1TJ: −50 ms. Traces are grouped by random walk direction and trial-averaged across multiple sessions (ALM, n = 12 sessions; M1TJ: n = 9 sessions). Same plotting conventions as in **b**, except inter-lick intervals are included. **e**. (Left) Scatter plot of lick n+1 contact angle 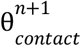 and the change from lick n to lick n+2 in port angle decoded from neural activity 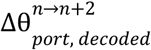 in ALM (n = 3245 control licks, 554 opto licks, 14 sessions). Licks without S1TJ optogenetic perturbation in grey, licks with S1TJ optogenetic perturbation in blue. Solid line indicates the line of best fit from linear regression. Sign of the lick contact angle 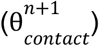 flipped for ease of visualization. Pearson’s correlation coefficient r shown in inset. *** p<0.001, one-sided paired shuffle test. (Right) Scatter plot of port angle decoded from neural activity 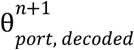 in ALM at lick n+1 and actual lick angle of lick n+2 ^*n*+2^. Same plotting conventions as in left panel. **f**. Similar scatter plot as **e**, but change in port angle decoded from neural activity 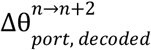 and port angle decoded from neural activity 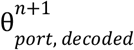 in M1TJ (n = 2723 control licks, 412 opto licks, 9 sessions).

We briefly inhibited S1TJ during the lick occurring after the random walk (lick n+1) in a subset of trials and recorded neurons in ALM and M1TJ (Extended Data Fig. 7a, b). While overall firing rates in the motor cortex for both putative pyramidal and fast-spiking neurons were reduced during S1TJ photoinhibition, there remained residual activity in a subset of neurons (Extended Data Fig. 3c, d, Extended Data Fig. 7a, b). Linear models of population activity without optogenetic perturbations were used to decode neural correlates in the presence or absence of optogenetic perturbations (Fig. 4). As expected, optogenetic perturbation of S1TJ impaired subsequent lick targeting (Fig. 4b) and reduced decoding accuracy of kinematics in the motor cortex population activity (Fig. 4c, d).

For the update of estimated lick port location, we investigated whether the neural correlate of lick port angle was updated by tactile feedback on a lick by lick basis (Fig. 4e, f; left). We measured the correlation between lick contact angle (θ_contact_) at lick n+1 and the change in lick port location decoded from population activity in the time interval between lick n and n+2 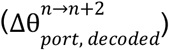. As expected, the correlation between lick contact angle and change in decoded lick port location in both ALM and M1TJ decreased significantly with S1TJ photoinhibition (p<0.001), indicating disruption in updating estimated lick port location.

Similarly, for the update of the intended lick angle, we investigated whether the neural correlate of current port location informed the angle of the subsequent lick (Fig. 4e, f; right). We measured the correlation between decoded lick port location at lick n+1 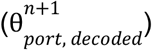 and the actual lick angle at lick n+2 (θ ^*n*+2^). As expected, the correlation between decoded lick port location and angle of the subsequent lick decreased significantly with S1TJ photoinhibition (p<0.001), indicating disruption in using the estimated lick port location to plan the angle of the next lick. In summary, optogenetic perturbation of somatosensory processing disrupts motor cortex estimation of lick target position and motor planning for the next lick angle.

### Persistent memory of target location in orofacial motor cortex

In the probabilistic sequence licking task, the goal is to keep track of the lick target location via somatosensory feedback. However, in the inter-trial interval when mice withhold licks, memory of lick target location needs to be maintained for seconds in the absence of somatosensory feedback from licks. Expert mice directed their initial licks in a given trial to the lick port position at the end of the previous trial (Fig. 5a), suggesting memory of target position persists during inter-trial intervals.

**Figure 5.**
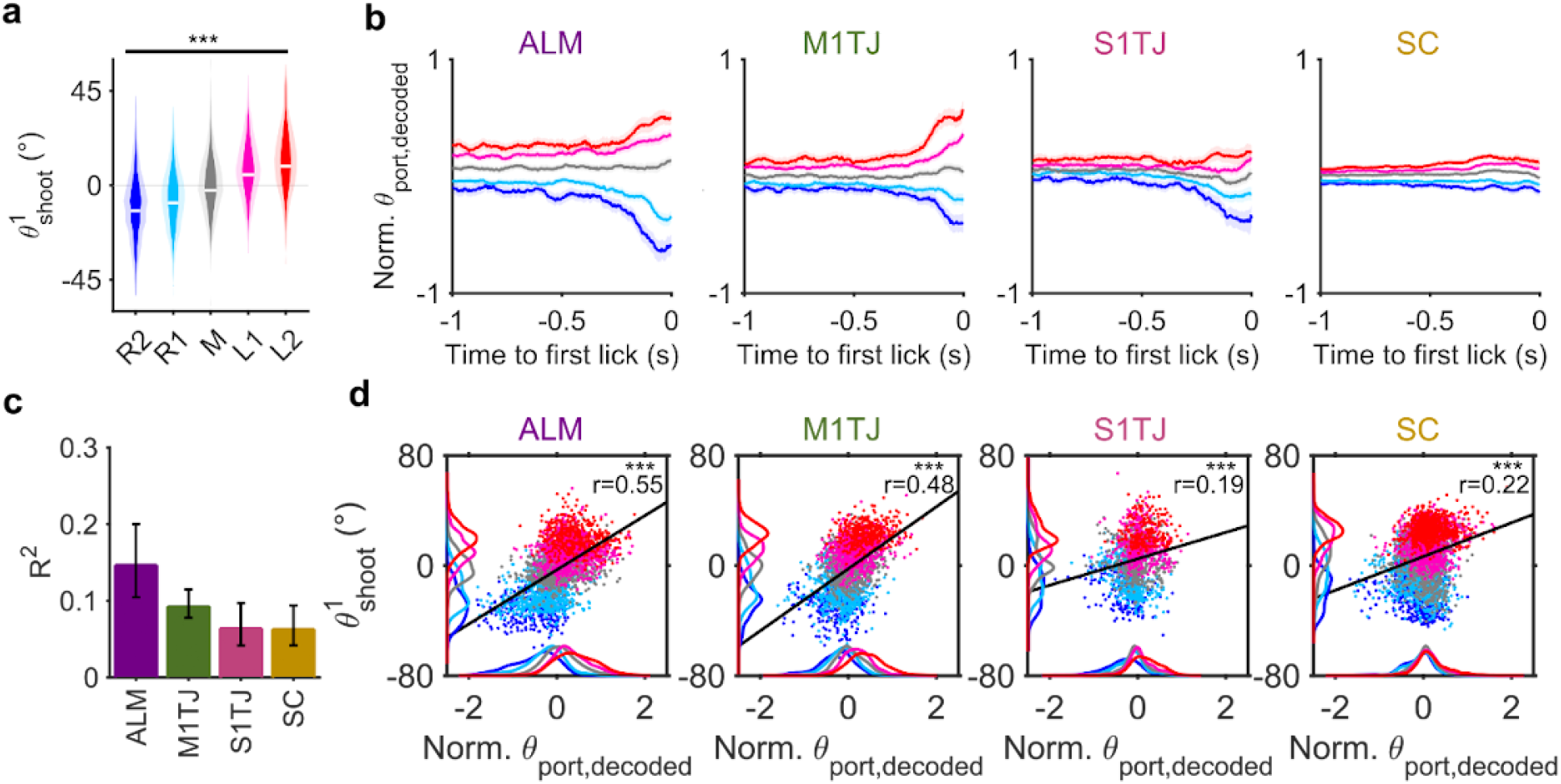
Persistent memory of lick target position is maintained in the orofacial motor cortex. **a**. Probability distributions of shooting angle for the first lick in the sequence 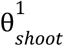 across lick port positions (n = 72 sessions, 7 mice; mean ± s.d.). White lines indicate medians of the probability distributions. *** p<0.001, Kruskal-Wallis test. **b**. Linear decoding of normalized lick port location from neural activity prior to first lick in the ALM, M1TJ, S1TJ, and SC (mean ± 99% bootstrap confidence interval; ALM, n = 12 sessions; M1TJ: n = 9 sessions; S1TJ, n = 6 sessions; SC n = 14 sessions). Lick port locations colored as R2, blue; R1, cyan, M, gray; L1, pink; L2, red. **c**. Goodness of fit of linear regression of lick port location from neural activity in ALM, M1TJ, S1TJ, SC (mean ± 95% bootstrap confidence interval). **d**. Scatter plots and kernel density estimations of decoded port location from neural activity prior to first lick in the ALM, M1TJ, S1TJ, and SC and shooting angle for the first lick in the sequence 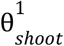 (ALM, n= 3063 trials, 12 sessions; M1TJ, n= 2731 trials, 9 sessions; S1TJ, n= 1743 trials, 6 sessions; SC, n= 5343 trials, 14 sessions). Lick port locations colored as R2, blue; R1, cyan, M, gray; L1, pink; L2, red. Black lines indicate the line of best fit from linear regression. Pearson’s correlation coefficient r shown in inset. *** p<0.001, one-sided shuffle test.

Given previous investigations into the maintenance of working memory in the frontal cortex in delayed-response tasks,^6,23,24^ we predicted that neural correlates for lick port position during the inter-trial interval would be strongly maintained in the orofacial cortex. Linear decoding of lick port positions showed large separation of decoded lick port positions in the population activity in ALM and M1TJ prior to first lick (Fig. 5b). In addition, we observed ramping activity, in which separation between the decoded positions widened over 200 ms prior to the first lick, similar to previous observations of ramping activity in the frontal cortex in delayed-response tasks. In comparison, separation of decoded lick port positions remained small with minimal ramping in SC, suggesting weaker encoding of target location in the absence of sensory feedback. In addition, on a trial-by-trial basis, the angle of the first lick of the trial correlated strongly with port location decoded from ALM and M1TJ activity during the inter-trial interval and weakly with port location decoded from S1TJ and SC activity (Fig. 5d). These observations indicate that the memory of lick target position is primarily maintained in the motor cortex.

## Discussion

Neural control of movement occurs across multiple levels and timescales.^25^ Guided by available sensory information in the environment, the brain can flexibly and rapidly assemble sequences of movements.^17,19,26,27^ In this study, we examined how a multiregional network across the motor cortex, somatosensory cortex, and superior colliculus can plan and direct licks to a target at each step of a probabilistic sequence based on immediate somatosensory feedback. Integration of sensory feedback, motor preparation, and execution from lick to lick depended on population activity in orofacial motor cortex and somatosensory cortex in the subsecond timescale. Our findings suggest that computations across cortical areas and superior colliculus generate an internal model of the movement target and continuously update the model based on sensory feedback.

Neural control of movement sequences flexibly constructed on-the-fly uses a dynamic and continual cycle of sensory feedback, motor planning, and motor execution. Our findings suggest that plans for current and future motor actions can be held concurrently in the motor system. In non-human primates performing a two-reach sequence,^12^ and in humans performing sequential finger taps under conflicting sensory and memory cues,^13^ rapid execution of sequential actions relied on the ability to independently plan for the next motor action while performing a disparate current action. Our work extends this idea of independent generation of motor actions at each step of a sequence to orofacial movements in mice.

Aiming licks in probabilistic sequences required activity in orofacial cortical areas, in contrast to one or two-step touch-guided lick aiming that did not require cortical activity.^20^ While targeting licks may in principle be distilled to stimulus-guided orienting based on a sensorimotor map in the superior colliculus, there are several features of the probabilistic sequence licking task that may necessitate cortical involvement. First, multi-step random walks in the sequence means that neural circuits execute a wide repertoire of motor sequences, which may require cortical networks for flexible sequence learning and execution, as previously described.^26,27^ Second, in contrast to single-step touch-guided lick aiming, in which the initial position of the lick port is held constant, the initial target position in probabilistic sequence licking is dynamic. This necessitates internal maintenance of information about target position, which is thought to rely on cortical processing.^11,23^ Finally, the role of cortical areas in driving licking movements can depend on the presence or absence of water rewards.^17,19,28^ As the water reward is given at the end of the entire probabilistic sequence, the licks in the sequence may be wholly dependent on cortical activity.

While motor control in probabilistic sequence licking depends on the orofacial cortex, it is by no means the sole region responsible for targeting licks. The orofacial cortex is a node in a multiregional circuit for tongue motor control,^24,25,29,30^ which includes top-down controllers, such as motor cortex^5,31^ and superior colliculus,^20,21^ and low-level controllers, such as brainstem central pattern generators (CPGs) for orofacial movements.^32–34^ How descending signals from the motor cortex, generated from structured population activity,^9,31,35,36^ interface with subcortical areas and brainstem CPGs in preparing and executing movements under varying task demands remains an open and interesting question for future studies.

## Methods

### Mice

All procedures were in accordance with protocols approved by the Johns Hopkins University Animal Care and Use Committee (protocols: MO18M187 and MO21M195). We obtained behavioral/neural data from 4 wild-type mice (1 female, 3 male; C57BL/6J, JAX #000664) and 19 double heterozygous VGAT-IRES-Cre/+; Ai32/+ (9 female, 10 male; obtained from crossing VGAT-IRES-Cre and Ai32; Jackson Labs: 028862; B6J.129S6(FVB)-Slc32a1^tm2(cre)Lowl^/MwarJ; Jackson Labs: 012569; B6;129S-Gt(ROSA)26Sor^tm32(CAG-COP4*H134R/EYFP)Hze^/J) at 3-10 months of age. Mice were housed in a room on a reverse light–dark cycle.

### Surgical procedures

#### Headpost implantation with clear skull preparation

For implantation of titanium headposts, mice were anesthetized with isoflurane (1-2% in O2; Surgivet). The scalp was excised, the periosteum was removed, and the skull was scored. A thin layer of Metabond (C & B Metabond) was applied. The headpost was affixed on top. To make the skull transparent, a thin layer of cyanoacrylate adhesive was then dropped over the entirety of the Metabond-coated skull and left to dry. A silicone elastomer (Kwik-Cast) was then applied over the surface to protect the transparent surface.

#### Craniotomies for electrophysiological recordings

Prior to planned silicon probe recording sessions, we made circular (~1 mm diameter) craniotomies over the target locations (ALM, AP 2.5, ML ±1.5 mm; M1TJ, AP 1.8, ML ±2.6 mm; S1TJ, AP 0.5, ML 3.7 mm; SC, AP −3.5, ML ±1.7 mm) that could accommodate multiple probe insertions. To facilitate ease of insertion, the dura was removed with sharp forceps. For the purpose of grounding the probes, a metallic screw was implanted in a smaller craniotomy over the right barrel cortex or visual cortex.

### Probabilistic sequence licking task

#### Task control

Mice performed sequential directed licking towards a moving lick port in a previously described behavioral rig.^17^ A stainless steel lick port was mounted on two-axis linear motors (LSM050B-T4 and LSM025B-T4, Zaber Technologies). The height of the lick port was adjusted via a manual linear stage (MT1/M, Thorlabs). The task was controlled by a Teensy 3.1 microcontroller and custom MATLAB (R2020b) and Arduino (1.8.13) software. Lick port contacts were detected by a custom lick detection circuit (Janelia Research Campus). The possible port positions were preset to 5 positions (L2, L1, Mid, R1, R2) of approximately equal distance from the tip of the jaw with spacing between positions corresponding to roughly half of tongue width (~15 degrees in arc).

During probabilistic sequence execution, the trial started with an auditory tone (0.1 s long, 15 kHz pure tone). Every third tongue-port contact triggered port movements one step to the right or left. At the rightmost or leftmost edge positions, the port moved one step towards the center during port movement. A built-in time delay of 20 ms between lick contact and triggered port movement, the long lick length required for lick contact (4 - 5 mm), and programmed arc movement of the lick port^17^ reduced the likelihood of lick port brushing against the tongue during lick port movement and revealing the direction of the lick port movement. After 3 random walks, the trial ended, and a water reward was dispensed by a water valve (LHDA0531415H, The Lee Company) after a short delay (250 ms). After an inter-trial interval drawn from a truncated exponential distribution (min 1.5s, max 3s, mean 2s), the next trial started with the initial lick port position the same as the last position in the previous trial.

To reduce the likelihood that mice used visual or auditory cues to locate the lick port, mice performed the task without visible light and under auditory masking conditions as previously described.^17^ To reduce the likelihood that mice used tactile feedback from facial fur and whiskers rather than the tongue surface to locate the lick port, the lower face and lowest row of whiskers were trimmed every 3-4 days. Previous work in the same behavior rig with hearing loss with earplugs and odor masking^17^ showed that mice directed licks at a moving lick port without relying on subtle auditory or olfactory cues.

#### Behavioral training

The behavioral training regime was similar to that previously described.^17^ Briefly, initial training sessions involved a short lick distance to port and a small range in angles between lick port positions. As the training progressed over 2-3 weeks and mice gained proficiency at modulating their lick angle (θ), we gradually increased both the lick distance to the port to ~ 4.5 mm and the distance between the successive lick port positions until the rightmost and leftmost positions differed by ~ 50 degrees. We considered mice to be proficient when they could successfully complete the probabilistic sequence in most trials in a rapid succession of licks (2 - 3 s long lick bouts) and complete a minimum of 200 trials in a 1 hr session.

#### Optogenetic stimulation

Optogenetic stimuli were applied using a pair of 470 nm LEDs (M595F2, Thorlabs) via an optic fiber (M98L01, Thorlabs) positioned over target areas with approximately 1 mm radius (ALM/M1TJ, AP 2.5, ML ±1.5 mm; S1TJ, AP 0.5, ML ±3.7 mm, 30 deg angle from vertical; S1BF, AP −1.5, ML ±3.5 mm). We used Wavesurfer (wavesurfer.janelia.org) to deliver 0.15 s (Fig. 2) or 2 s (Extended Data Fig. 2) stimuli consisting of 40 Hz sinusoidal waveforms, with a 25 or 100 ms ramp down period at the end. In 30% of trials in a given session, the stimuli were triggered with a 120 ms delay following the tongue-port contact prior to the random walk (lick n) and randomized to occur once in a given trial following any one of the lick contacts prior to the random walk (lick n or n’ or n”) in the sequence. In photoinhibition experiments over a transparent skull, the LED power at the tip was set to peak 8 mW, average 4 mW. In photoinhibition experiments over craniotomies, the power was set to peak 4 mW, average 2 mW. We supplied randomly flashing masking light of similar wavelength to prevent anticipation of optogenetic stimuli.

### Quantification of behavioral variables

#### High speed videography

We captured high speed video of the mouse performing the task at 400 frames per second. Individual frames were hardware-triggered and captured bottom and side views of the mouse’s face, which was illuminated with an 850 nm LED (LED850-66-60, Roithner LaserTechnik). Videos were acquired through a telecentric lens (0.25X, Edmund Optics), using a PhotonFocus DR1-D1312-200-G2-8 camera and Streampix 7 software (Norpix). Videos for each trial were captured separately, up to a maximum length of 20 seconds.Trials of duration exceeding 20 seconds were discarded from analyses.

#### Tongue kinematics

We used DeepLabCut^15^ to fine-tune ImageNet-trained ResNet-50 convolutional neural networks (CNN) to annotate the tongue tip, tongue base, and jaw tip in each frame. The tip of the lick port was marked manually for each of the five lick port positions. To smooth out frame-to-frame jitter in the CNN outputs, we applied a median filter of 5 frames. Distance between tongue tip and base was used to determine frames in which the tongue was visible (tip-base distance greater than 5 pixels or 20% of maximum, whichever was greater). Video frames in which the tip-base distance exceeded the threshold for more than 5 consecutive frames were labelled as frames containing licks. The reference point was determined by computing the median position of the jaw tip during periods without licking movements across all trials in a given session.

Lick length (L) was defined as the distance between the reference point and the tongue tip. Lick angle (θ) was defined as the angle of the vector connecting the reference point and the tongue tip. Port angle (θ_port_) was defined as the angle of the vector connecting the reference point and the lick port tip. By convention, negative values denoted angles to the right of the vertical (left-ward licking or left port positions), while positive values denoted angles to the left.

Lick contact angle (θ_cortact_) was computed by subtracting the port angle from lick angle (θ_cortact_ = θ - θ_port_). For Fig. 4, we flipped the sign of the lick contact angle (−θ_cortact_ = θ_port_ - θ) on the x-axis for ease of visualization in correlating lick contact angle with neural correlates. Determination of port angle (θ_port_) and lick contact angle (θ_cortact_) was restricted to frames with tongue-port contact. In Fig. 2, Extended Data Fig 1, 5, the lick contact angle was averaged over the duration of each individual contact to yield a single scalar value per touch event. As previously described,^17^ we used θ_shoot_, defined as the θ when L reached 0.84 maximal L (Lmax), to succinctly depict the lick angle as a single scalar value per lick. In Extended Data Fig. 4, error in lick angle (θ_error_) was computed identically as lick contact angle (θ_cortact_) for licks with touch events. For licks without touch events, error in lick angle (θ_error_) was computed by finding the point in the lick trajectory closest to the lick port and computing the difference in angle between the lick vector and lick port vector at the closest point.

#### Lick grouping, alignment, and standardization

In quantifying behavior, we pooled together licks by their order relative to the random walk, grouping licks (n’, n”, n’’’) together as lick n and similarly for licks (n’+1, n”+1, n’’’+1) as lick n+1 and so forth, and did not distinguish between their absolute order within a trial.

As previously described,^17^ prior to analyses that were sensitive to variable lick rates (Fig. 3, 4), we linearly warped inter-lick intervals within each lick bout to a constant value of 0.154 s, corresponding to overall mean lick rate of 6.5 Hz. Original time series, including spike rates, were obtained before warping inter-lick intervals. After alignment, the behavioural and neural time series were resampled uniformly at 400 samples per second.

Similarly, a subset of analyses were sensitive to different ranges of lick angles, lengths, and port angles due to differences in head-fixation, camera view angles, or idiosyncratic licking preferences (Fig. 3, 4). Thus, for each session, we standardized the range of lick angles by zero-meaning the shooting angle (θ_shoot_) of licks to the middle lick port position and scaling the difference in shooting angle between licks to the rightmost and leftmost positions to 50 degrees. We zero-meaned and scaled the port angle (θ_port_) by the same normalization factor.

#### Lick cycle timing

We quantified the variability in the timing of optogenetic perturbation relative to the lick cycle (Extended Data Fig. 4) with the metric Δt, the difference in time between onset of optogenetic perturbation and the mid-lick time. Mid-lick time was defined as the time midpoint of video frames in which the tongue of a given lick was visible. Mid-lick time was used as the default timestamp of a given lick. To quantify kinematics as a function of variability in photoinhibition timing, each lick during and after photoinhibition was binned by its associated Δt with bin widths of 20 ms (L) or 40 ms (θ_error_). Bins with an insufficient number of licks (<5 licks or <0.5% of total lick count) were ignored. For control licks without photoinhibition, the metric Δt was computed by computing the difference in time between onset of a fictitious optogenetic stimulus (occuring 120 ms following lick n) and the mid-lick time.

### Electrophysiology and spike sorting

#### Neural recordings

We used single-shank 384-channel Neuropixel 1.0 silicon probes to extracellularly record neurons while mice performed the task. A small subset of recordings used four-shank Neuropixels 2.0 probes. The targeted regions were the following; ALM, AP 2.5, ML ±1.5 mm, depth 1350-1450 µm; M1TJ, AP 1.8, ML ±2.6 mm, depth 1350-1450 µm; S1TJ, AP 0.5, ML 3.7 mm, depth 1250-1350 µm, 40 degree angle from vertical; SC, AP −3.5, ML ±1.7 mm, depth 2600 µm. Prior to probe insertion, the probe was coated with lipophilic dye (Vybrant CM-DiI, ThermoFisher). Probe insertion was stabilized with agarose gel. Data were acquired at ~30 KHz using a PXIe chassis (PXIe-1071, National Instruments) equipped with control and acquisition modules (IMEC Neuropixels PXIe control system and PXIe-8381, PXIe-6341, National Instruments), and SpikeGLX software (billkarsh.github.io/SpikeGLX).

#### Spike sorting and pre-processing

Voltage traces were processed using the Ecephys spike sorting pipeline (github.com/jenniferColonell/ecephys_spike_sorting) with Kilosort 2.5.^37^ Clusters were selected as putative single units on the following quality metrics:

I. Inter-spike interval violations (a measure of contamination by other units) < 0.5
II. Amplitude cutoff (a measure of missing spikes) < 0.1
III. Presence ratio (a measure of completeness of the data) > 0.8

Putative single units were then manually assessed via inspection in Phy (github.com/cortex-lab/phy) for manual merging or splitting.

For cortical recordings, we classified units as putative pyramidal neurons if the width of the average spike waveform (defined as time from trough to peak) was greater than 0.5 ms, and as putative fast-spiking interneurons if shorter than 0.4 ms.

#### Assessing false discovery rate

Using the analytical model previously described (github.com/economolab/DCISIv),^22^ we assessed the cross-contamination rate in our electrophysiology dataset. Using the inter-spike interval violation rates of simultaneously recorded single units, we estimated the population-level false discovery rate for each recording session, using a homogenous firing rate model.

### Histology

After completion of all neural recording experiments, we perfused mice transcardially with 4% paraformaldehyde in phosphate buffered saline, and extracted the brains. Following overnight post-fixation, we obtained 150 μm coronal slices (Pelco easiSlicer, Ted Pella) and captured widefield epifluorescence images (AxioZoom16, Zeiss). Slice images were registered to the Allen CCF. Probe tracks reconstructed using SHARP-Track (github.com/cortex-lab/allenCCF), and used to confirm placement of probes for a given brain region.

### General data analysis

We used MATLAB (R2021a, Mathworks) for data pre-processing, and Python (3.8) and associated libraries for analyses, except where otherwise noted. We did not pre-determine any sample sizes. In a subset of analyses involving trials pooled across sessions and mice, we used hierarchical bootstrap resampling as follows: to generate bootstrap samples, we randomly sampled (with replacement at each level) mice, then sessions from the sampled mice, and finally trials from the sampled sessions. We computed the relevant test statistic for each bootstrap sample. Reported confidence intervals were obtained from percentile values of these resampled test statistics.

In analyses comparing an observed dataset to a shuffled dataset, we used hierarchical shuffling such that relevant quantities (e.g. lick pairs) were shuffled not globally across animals and sessions, but locally within individual sessions, and then used to compute relevant test statistics. In analyses capturing the effect of optogenetic stimulation, we used a paired shuffling strategy such that group labels (control vs optogenetics) were shuffled locally within individual sessions and differences in relevant quantities between shuffled groups were computed as the relevant test statistics.

### Firing rates and peristimulus time histograms

To compute peristimulus time histograms (PSTHs) in response to optogenetic stimulation or random walks, we computed the firing rate of each unit for each trial in 2.5 ms bins, over which we applied a Gaussian smoothing kernel in time, and aligned each trial to the relevant time point (optogenetic stimulation onset or lick n). Mean firing rate (PSTH) and standard error was computed by averaging over relevant trials. For normalized PSTHs, we normalized the mean firing rate of a unit by its maximum mean firing rate. Units with firing rates below 1 Hz in trials without optogenetic stimulation were omitted from the normalized PSTH plot.

### Linear regression and decoding

#### Preprocessing

For linear regression and decoding, sessions with insufficient unit counts (<15 single units) or insufficient numbers of trials (<150 trials) or probes located in incorrect regions were excluded from analysis. In time lag analysis, a range of time lags (−150 to 150 ms) were added to spike times prior to computing firing rates. For each session, firing rates of simultaneously recorded units were ‘soft’ normalized with a factor of the maximum firing rate + 5 Hz. The time bins used for decoding behavioral parameters (θ_contact_, θ_port_, θ) following a random walk were t = [−0.1, 0.4] (s) relative to time of lick n (midpoint of visible tongue frames). The time bins used for decoding normalized θ_port_ were t = [−1, 0] (s) relative to time of first lick of a trial. Normalized port angle θ_port_ was derived by setting the range of port location (R2, R1, M, L1, L2) to (−2, −1, 0, 1, 2).

#### Fitting linear models

We used firing rates during trials without optogenetic stimulation to fit linear models. For each session, we used linear regression with elastic net regularization (‘lasso’ in Matlab) to fit each individual behavioral parameter (θ_contact_, θ_port_, θ, normalized θ_port_) as a linear weighted sum of simultaneously recorded firing rates. Elastic net regularization in Matlab uses mixing parameter ɑ for relative weighting of L1 and L2 norms and overall regularization parameter λ. Mixing parameter α was chosen on a session-by-session basis from among candidate values {0.001, 0.01, 0.1, 0.2, 0.5, 1} as the value that yielded the minimum 5-fold cross-validated mean square error (MSE). Using the optimal mixing parameter α, we used the default Matlab set of regularization parameters λ, a geometric sequence of 100 λ values with largest λ_max_ giving a non-null model and 1e-4 ratio between values. With 10-fold cross-validation, we chose the optimal λ value that minimized MSE. We measured decoding performance of linear models by computing the 10-fold cross-validated coefficient of determination (R^2^). Using an alternative λ value (largest λ value such the MSE within one standard error of the minimum MSE) did not affect decoding performance significantly. For a subset of analyses, we averaged the decoded neural correlates for each trial/lick over relevant time bins for n, n+1, n+2 lick durations or for pre-lick periods [−0.2, −0.05] (s) relative to the time of first lick. To obtain an explicit null model and its corresponding coefficient of determination, we shuffled each unit’s spike rate in time and repeated the above fitting procedure.

## Acknowledgments

We thank Isis Wyche, Yi-Ting Chang, Reza Shadmehr, Yujin Han and Mike Economo for suggestions and discussion; Yujin Han and Mike Economo for comments on the manuscript. This work was supported by NIH grants T32GM136577, 1F31DE033256 and 1U19NS137920-01.

## Author contributions

Conceptualization, J.J.K. and D.H.O.; investigation, J.J.K.; methodology, J.J.K. and M.D.; formal analysis, J.J.K.; writing – original draft, J.J.K.; writing – review & editing, J.J.K., M.D. and D.H.O.; funding acquisition, J.J.K. and D.H.O.; supervision, D.H.O.

---

**Extended Data Figure 1.**
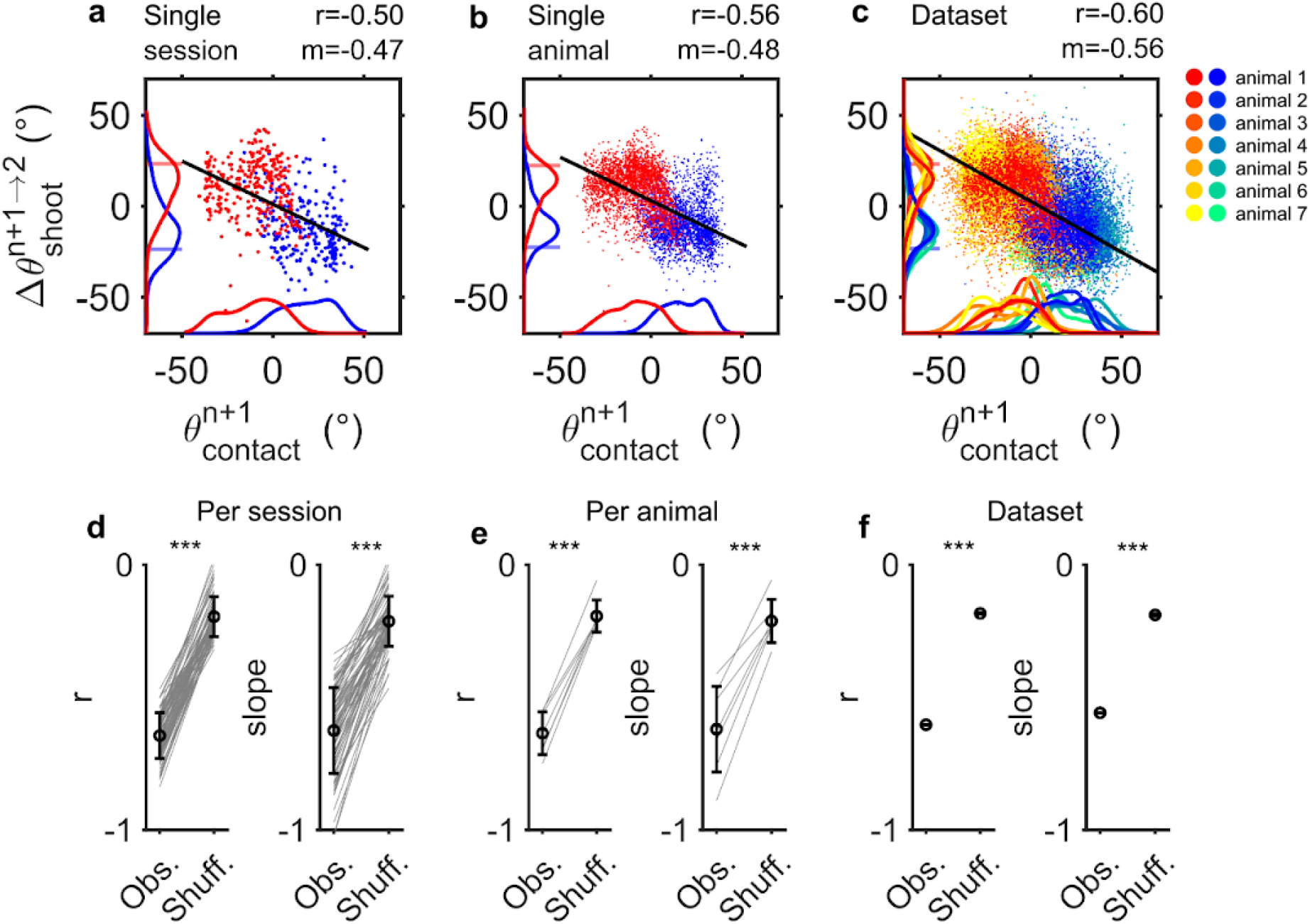
Lick angle adjustment based on contact angle occurs for each session and animal. **a**. Scatter plot and kernel density estimation of lick n+1 contact angle 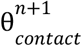 and subsequent change in lick shooting angle.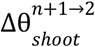 for a single session. Same plotting conventions as Fig. 1e **b**. Same as **a** for a single animal. **c**. Scatter plot and kernel density estimation of lick n+1 contact angle 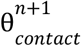 and subsequent change in lick shooting angle 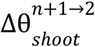, colored by the identity of the animal. **d**. Pearson correlation r and slope of linear regression of lick n+1 contact angle 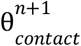 and subsequent change in lick shooting angle 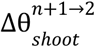 Δθ per individual session (n = 72 session) compared across observed and shuffled dataset. Individual sessions in gray. Mean ± s.d. In black. *** p<0.001, one-sided shuffle test. **e**, Same as **d** for per animal (n = 7 animals). Individual animals in gray. **f**. Pearson correlation r and slope of linear regression of lick n+1 contact angle 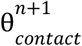 and subsequent change in lick shooting angle 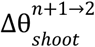 Δθ compared across observed and shuffled dataset. *** p<0.001, one-sided shuffle test.

**Extended Data Figure 2.**
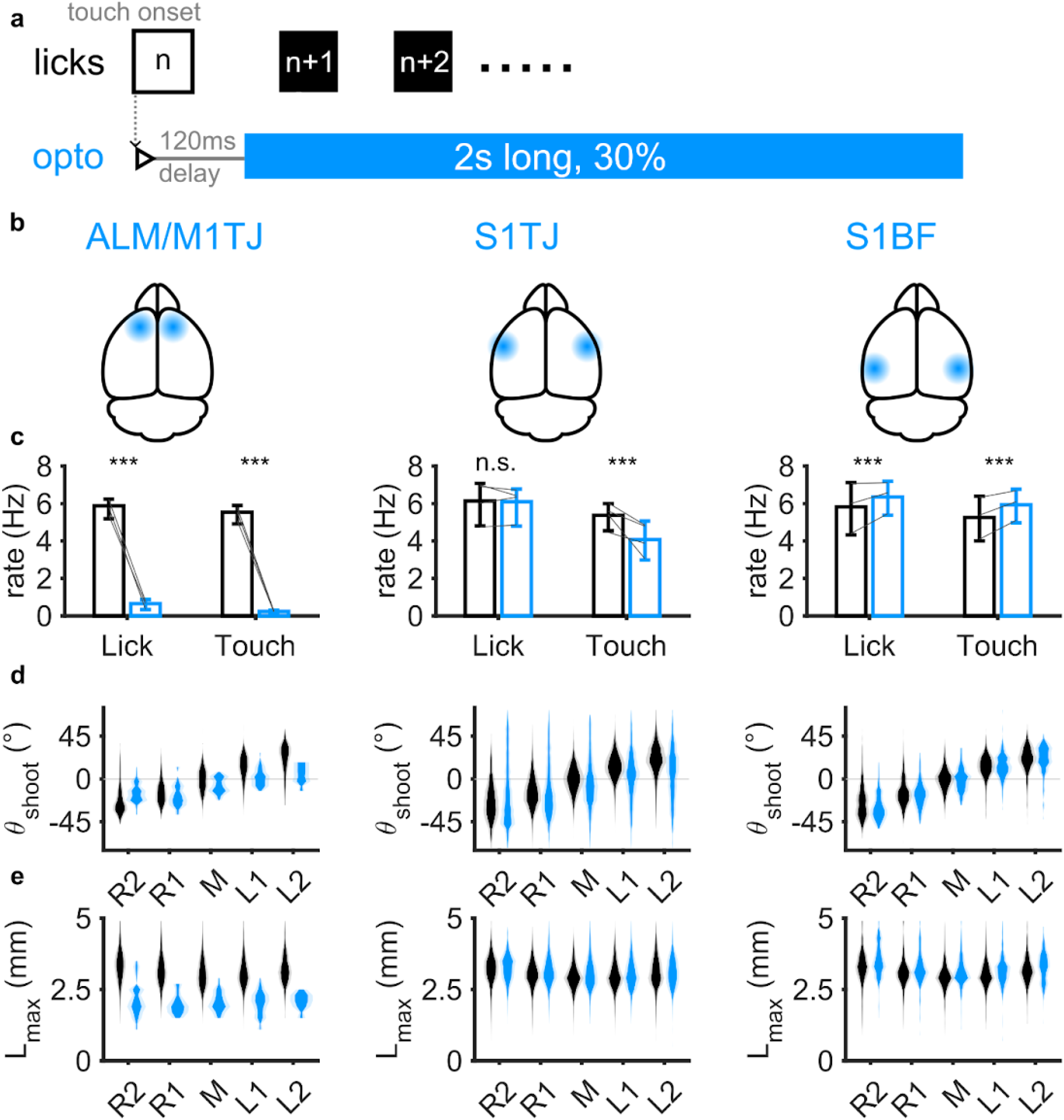
Bilateral photoinhibition of orofacial motor cortex reduces lick rate and length, and bilateral photoinhibition of orofacial somatosensory cortex reduces accuracy in lick angles. **a**. Diagram of closed loop optogenetic perturbation of lick n+1. Photoinhibition (40Hz sinusoid with ramp-down) occurs randomly in 30% of trials following a 120 ms delay after touch onset of lick n and lasts for 2 s. **b**. Schematic of cortical regions targeted for photoinhibition; ALM, anterolateral motor cortex; M1TJ, tongue-jaw primary motor cortex; S1TJ, tongue-jaw primary somatosensory cortex; S1BF, barrel cortex. **c**. Changes in the rate of licks and touch when inhibiting each of the three cortical areas (columns). Bar plots show mean ± 99% hierarchical bootstrap confidence interval with blue color for licks during optogenetic perturbation and black color for licks without optogenetic perturbation. Rate of lick and touch for each mouse shown in grey (ALM/M1TJ, n = 5 sessions, 3 mice; S1TJ: n = 16 sessions, 4 mice; S1BF, n = 3 sessions, 3 mice). *** p<0.001, two-sided paired shuffle test. **c**. Probability distributions of shooting angle θ_shoot_ for licks with (blue) or without (black) optogenetic perturbation for each of the three cortical areas (columns) (mean ± s.d). **d**. Probability distributions of maximum length L_max_ for licks with (blue) or without (black) optogenetic perturbation for each of the three cortical areas (columns) (mean ± s.d).

**Extended Data Figure 3.**
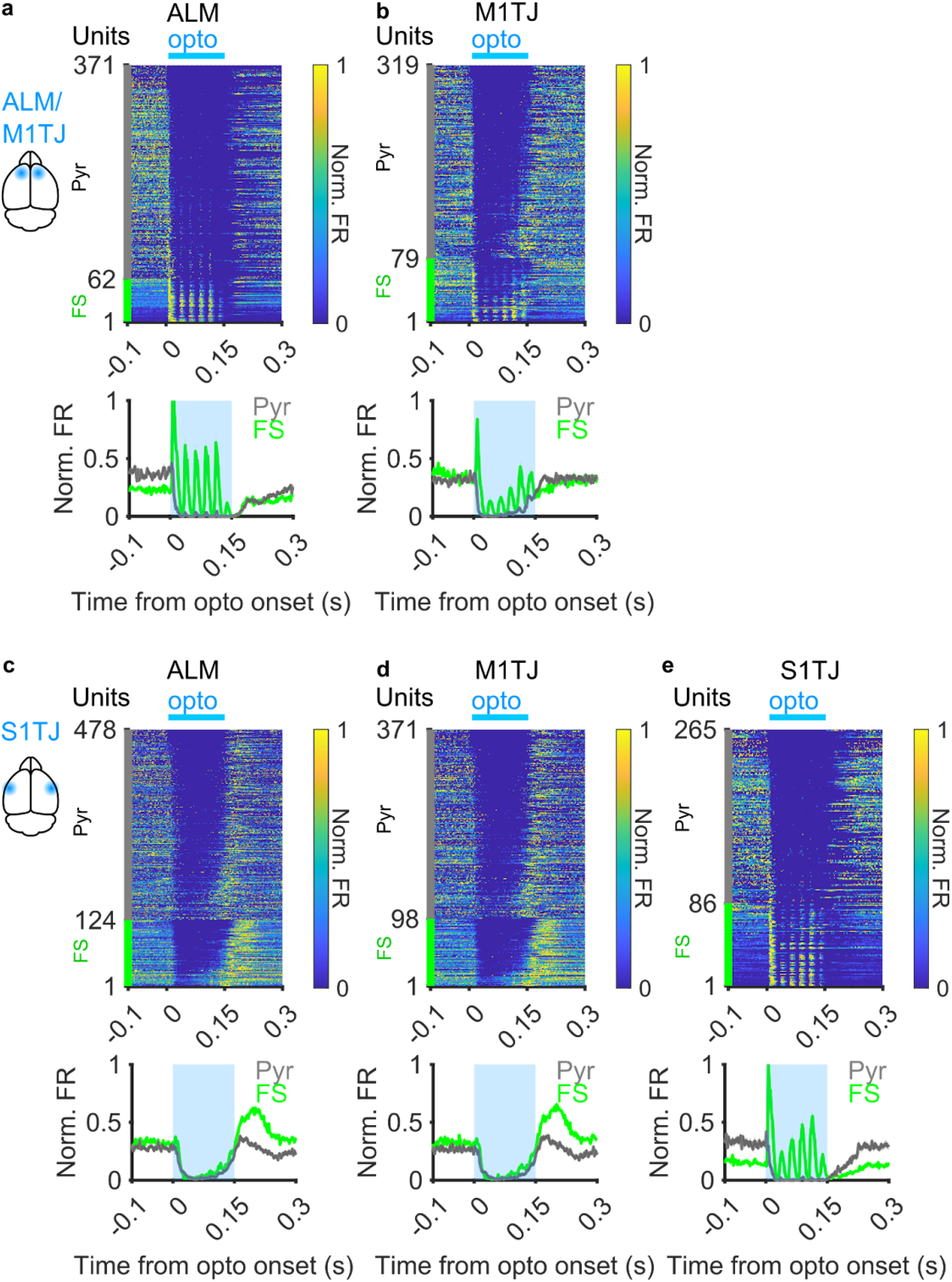
Electrophysiological validation of photoinhibition of orofacial cortical areas. **a-b**. (Top) Normalized trial-mean firing rates of neurons in cortical areas recorded (columns) during ALM/M1TJ photoinhibition. Grey and green indicate putative pyramidal (Pyr) and fast-spiking interneurons (FS), respectively. (Bottom) Average normalized firing rate across all recorded units in a given cortical area during ALM/M1TJ photoinhibition. **c-e**. Same as **a-b**, but during S1TJ photoinhibition.

**Extended Data Figure 4.**
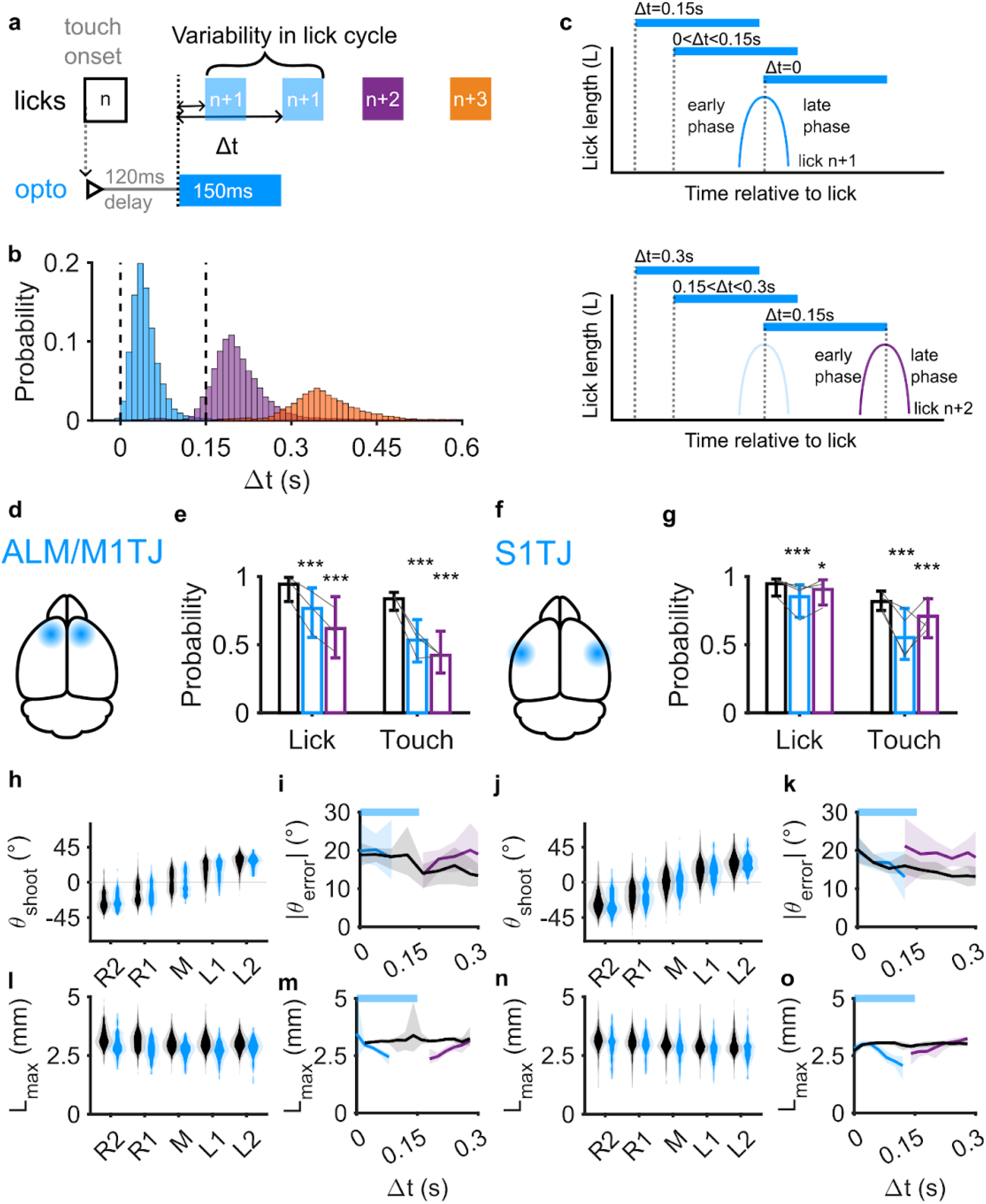
Variability in photoinhibition timing in a lick cycle does not affect lick angle, but affects lick length. **a**. Diagram of variability in timing of optogenetic perturbation relative to lick n+1.Given the variability in lick cycle duration (~6-7 Hz), the time delay Δt between the onset of optogenetic stimuli (triggered with fixed delay of 120 ms after lick n) and time of lick n+1, n+2, and n+3 can vary. **b**. Probability distribution of time delay Δt between the onset of optogenetic stimuli and time of lick (n = 72 sessions, 7 mice). Lick n+1 as blue, n+2 as purple, n+3 as orange. Dotted lines indicate onset and offset of optogenetic stimuli. **c**. Schematic of varying time delay Δt relative to lick n+1 (top) or lick n+2 (bottom). For lick n+1 (top), increasing Δt equals greater overlap between optogenetic stimuli and the early phase of the lick cycle. For lick n+2, decreasing Δt equals greater overlap between optogenetic stimuli and the early phase of the lick cycle. **d**. Schematic of motor cortex targeted for photoinhibition; ALM, anterolateral motor cortex; M1TJ, tongue-jaw primary motor cortex. **e**. Changes in the probability of licks and touch when photoinhibiting ALM/M1TJ. Bar plots show mean ± 99% hierarchical bootstrap confidence intervals with blue color for lick n+1 during optogenetic perturbation, purple color for lick n+2 after optogenetic perturbation, and black color for lick n+1 without optogenetic perturbation (ALM/M1TJ, n = 10 sessions, 3 mice). * p<0.05, *** p<0.001, two-sided paired shuffle test. **f**. Schematic of somatosensory cortex targeted for photoinhibition; S1TJ, tongue-jaw primary somatosensory cortex. **g**. Same as **d**, but photoinhibiting S1TJ (S1TJ, n = 17 sessions, 4 mice). **h**. Probability distributions of shooting angle θ_shoot_ for licks with (blue) or without (black) optogenetic perturbation in ALM/M1TJ (mean ± s.d). **i**. Magnitude of difference between lick angle and port angle |θ_shoot_| with respect to variability in ALM/M1TJ photoinhibition timing Δt (mean ± 95% hierarchical bootstrap confidence interval). Colors same as **d**. **j, k**. Same as **g, h**, but with optogenetic perturbation in S1TJ. **l**. Probability distributions of maximum length L_max_ for licks with (blue) or without (black) optogenetic perturbation in ALM/M1TJ (mean ± s.d). **m**. Maximum length L_max_ with respect to variability in ALM/M1TJ photoinhibition timing Δt (mean ± 95% hierarchical bootstrap confidence interval). Colors same as **d**. **n, o**. Same as **k, l**, but with optogenetic perturbation in S1TJ.

**Extended Data Figure 5.**
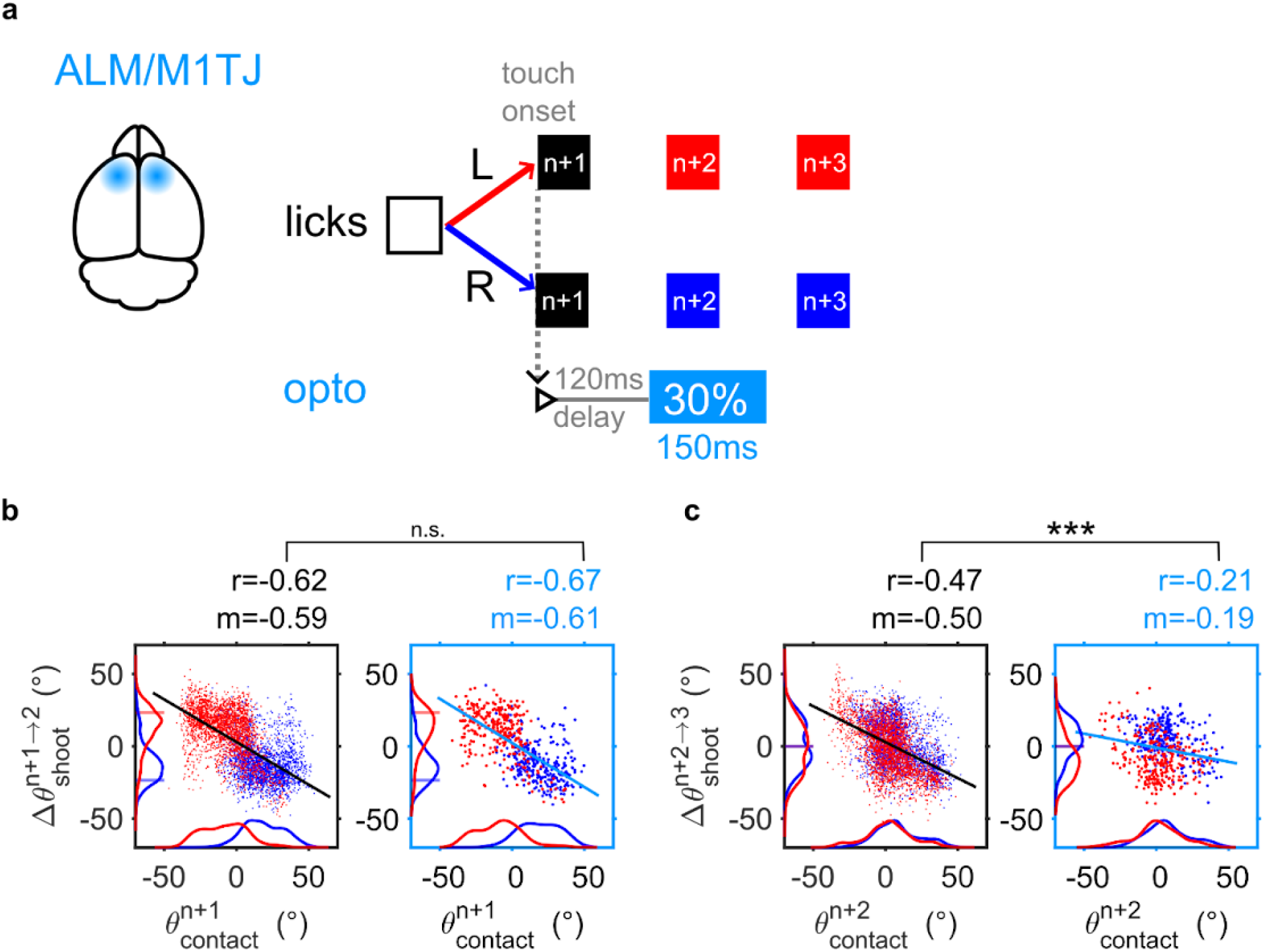
Planning in the motor cortex for the subsequent lick angle occurs in the current lick cycle. **a**. Diagram of closed loop ALM/M1TJ optogenetic perturbation of lick n+2. Photoinhibition (40Hz sinusoid with ramp-down) occurs randomly in 30% of trials following a 120 ms delay after touch onset of lick n+1 and lasts for 150 ms (Methods). **b**. Scatter plots and kernel density estimations of lick n+1 contact angle 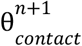 with or without ALM/M1TJ photoinhibition of lick n+2 (right, left) and subsequent change in lick shooting angle from lick n+1 to n+2 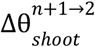 (n= 4852 control lick pairs, 612 opto lick pairs, 7 sessions, 3 mice). Plotting conventions same as in Fig. 2. **c**. Similar plot as **b**, but for lick n+2 contact angle 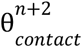 and subsequent change in lick shooting angle from lick n+2 to n+3 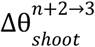

**Extended Data Figure 6.**
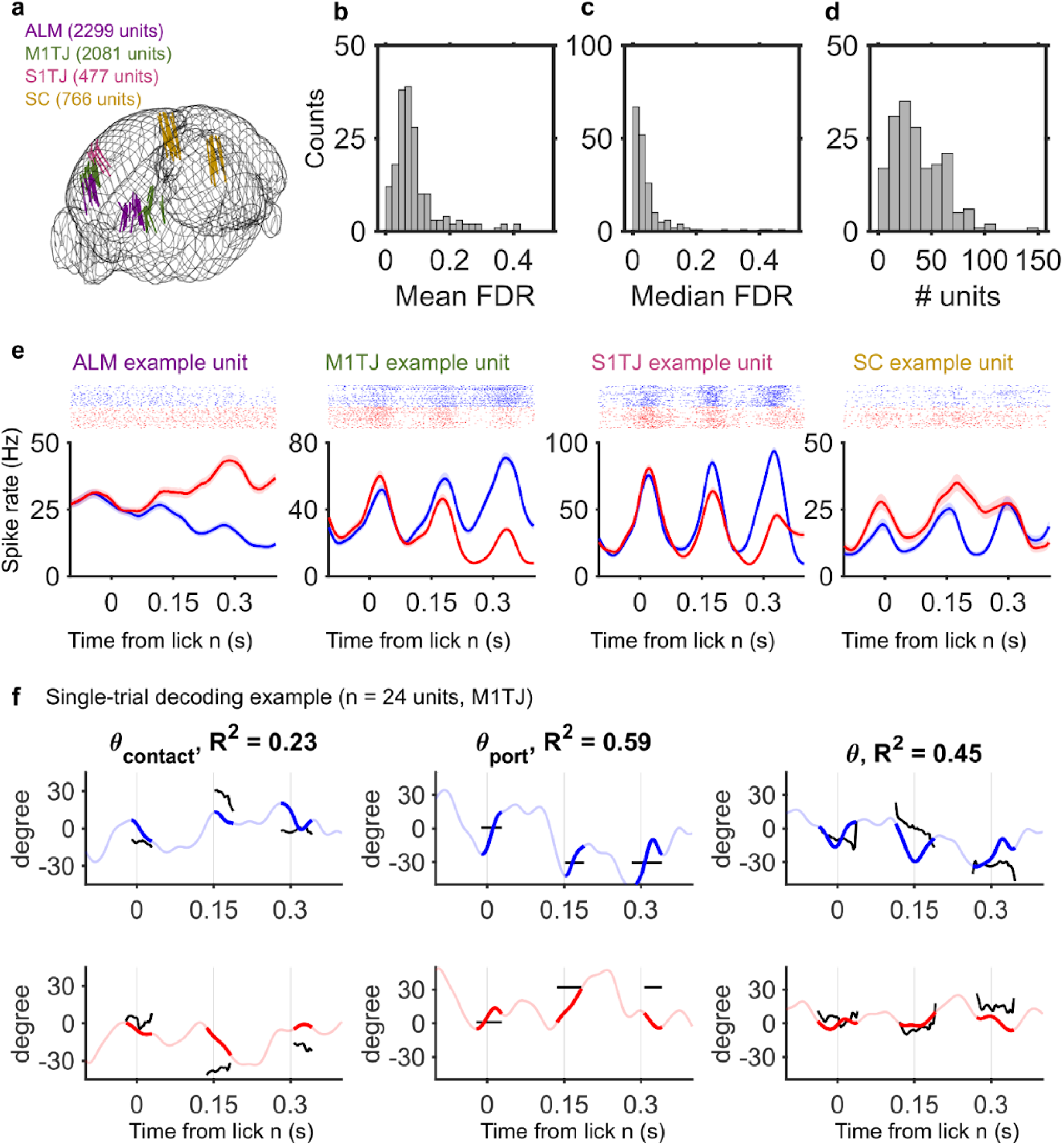
Characterization of electrophysiological recordings. **a**. Probe tracks for electrophysiological recordings in ALM, M1TJ, S1TJ, and SC. A subset of SC recordings had concurrent V1 unit recordings. **b-d**. Distributions of estimated mean false discovery rate, estimated median false discovery rate, and number of simultaneously recorded single units per session (n=182 recordings). **e**. Example single unit responses from ALM, M1TJ, S1TJ and SC recordings. (Top) Spike rasters from a subset of 50 trials in a given session. (Bottom) Peristimulus time histograms (PSTHs, mean ± s.e.) of single units aligned to time of lick n. Right and left random walks are color coded as blue and red, respectively. **f**. Single-trial decoding of behavioral variables (rows; black traces) from 24 simultaneously recorded M1TJ units for right (top, blue) and left (bottom, red) random walks. Solid lines indicate durations of visible tongue, shaded lines indicate inter-lick intervals. Goodness-of-fit (R^2^) for linear model for the given recording shown in title.

**Extended Data Figure 7.**
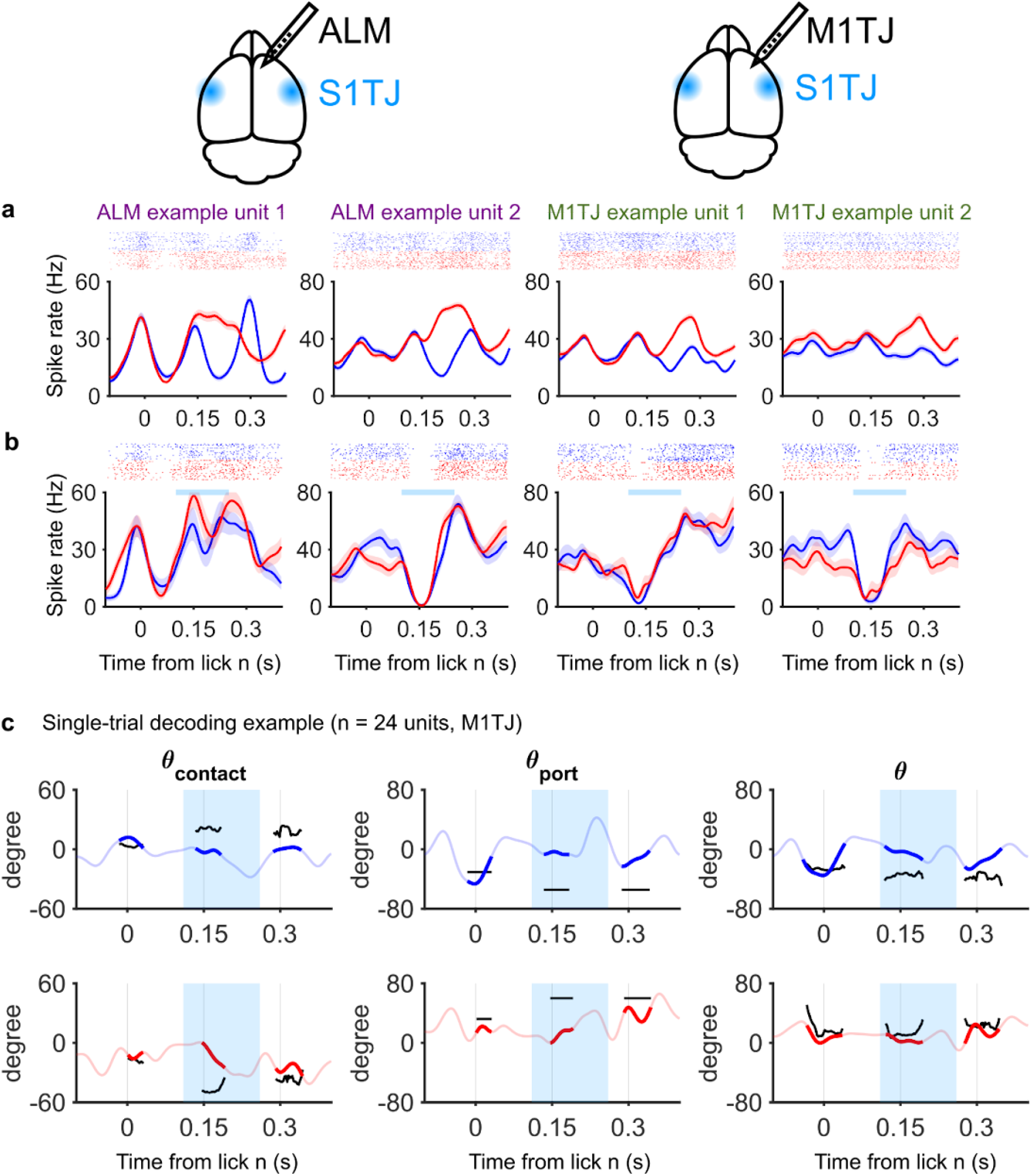
Example unit responses and decoding during S1TJ inhibition. **a**,**b**. Example single unit responses from ALM and M1TJ recordings without (**a**) or with (**b**) S1TJ photoinhibition. Duration of optogenetic perturbation in S1TJ indicated with cyan bar. Same plotting conventions as Extended Data Fig. 6. **c**. Single-trial decoding of behavioral variables (rows; black traces) from 24 simultaneously recorded M1TJ units with S1TJ photoinhibition for right (top, blue) and left (bottom, red) random walks. Same plotting conventions as Extended Data Fig. 6.

